# Clonal dynamics of monozygotic twinning in early human embryogenesis

**DOI:** 10.1101/2025.10.05.680569

**Authors:** Christopher Jongsoo Yoon, Chang Hyun Nam, Seung Mi Lee, Eun Saem Choi, Ji Hye Bae, Haemin Kim, Young Mi Jung, Joonoh Lim, Ryul Kim, Catherine Derom, Eline Meireson, Steven Weyers, Jung Woo Park, Junehawk Lee, Joohon Sung, Obi L. Griffith, Malachi Griffith, Jong Kwan Jun, Young Seok Ju

**Affiliations:** Graduate School of Medical Science and Engineering, Korea Advanced Institute of Science and Technology, Daejeon, Republic of Korea; Department of Medicine, Washington University School of Medicine, St. Louis, MO, USA; Department of Medicine, Johns Hopkins University School of Medicine, Baltimore, MD, USA; Department of Obstetrics and Gynecology, Seoul National University College of Medicine, Seoul, Republic of Korea; Inocras Inc., 6330 Nancy Ridge Drive, San Diego, CA, USA; Department of Human Structure and Repair, Ghent University, Ghent, Belgium; Department of Obstetrics and Gynecology, Ghent University Hospital, Ghent, Belgium; Korea Institute of Science and Technology Information (KISTI), Daejeon, Republic of Korea; Department of Epidemiology, School of Public Health, Seoul National University, Seoul, Republic of Korea

## Abstract

Monozygotic twins are derived from the split of a single zygote early in embryogenesis. Although it was hypothesized that the timing of twining is overall associated with fetal membrane configuration of twins, i.e., chorionicity and amnionicity, our understanding of early embryonic clonal dynamics underlying human twinning is limited. Here we explored the segregations of early embryonic lineages in 7 dichorionic diamniotic (**DCDA**), 7 monochorionic diamniotic (**MCDA**), 8 monochorionic monoamniotic (**MCMA**) monozygotic twins, and 1 dichorionic triamniotic (**DCTA**) monozygotic triplets, using post-zygotic early embryonic mutations (**EEMs**) as endogenous lineage barcodes. Patterns of the early lineage distributions among monozygotic twins revealed three apparent clonal categories, referred to as para-identical, sub-identical, and full-identical twins, which largely correlated with the amnionicity of the twins. Rather, despite conventional wisdom, chorionicity was not substantially associated with early clonal compositions, but with blood exchanges *in utero*. In sub-identical twins, where one co-twin was clonally a part of the other, our data suggested that the foundation of the latter co-twin was established after acquisition of a median of 6 additional post-zygotic mutations (range: 2–13), corresponding to ∼5 early cell divisions. Additional *in-depth* analysis on the matched placenta from an MCDA twin suggested that separation of two co-twins can precede the separation of the placenta and embryonic proper, and a single chorion can be formed even with multiclonal origin. Our findings provide insights into the clonal dynamics, twinning processes, and cell fate decisions in early human embryogenesis.

## INTRODUCTION

Multicellular organisms derived from a single zygote are composed of mosaic genomes resulting from somatic mutations introduced during early embryogenesis (Stratton et al. 2009; Ju et al. 2017). Monozygotic twins are a unique set of two individuals that are derived from a single zygote, and like other multicellular organisms, the cells in a monozygotic twin represent a collection of mosaic genomes. While often referred to as ‘identical twins’, due to the varying mutational lineage history of cells in each individual at the time of twinning, the genomic composition of each monozygotic twin sibling (hereafter referred to as co-twin) could not be perfectly identical (Dal et al. 2014; Jonsson et al. 2021). Although tracing early lineages of monozygotic twins, in principle, would provide deep insights into the twinning timing and processes, our knowledge is limited due, in part, to ethical constraints on experimental observations of human embryos.

Seven decades ago, Dr. George Corner proposed that the arrangement of fetal membranes (chorion and amnion) reflects the timing of embryonic developmental events, based on his studies of miscarried human embryos (Corner 1955)(**Fig. 1a**). Here, if the twinning process occurs very early (within the first 3 days after fertilization), two twin embryos form separate chorions and amnions, resulting in a dichorionic diamniotic (**DCDA**) twin. If embryo divisions occur later, such as on days 4-8 and days 8-13, twin embryos share a chorion but have separate amnions (monochorionic diamniotic; **MCDA**), and share both the chorion and the amnion (monochorionic monoamniotic; **MCMA**), respectively. If divisions occur after 13 days, conjoined twins with incomplete embryo separation can be generated. Despite the lack of direct experimental proof, the simple and intuitive hypothesis was soon incorporated into many reviews and textbooks (Herranz 2015; Hall 2003; McNamara et al. 2016).

**Figure 1.**
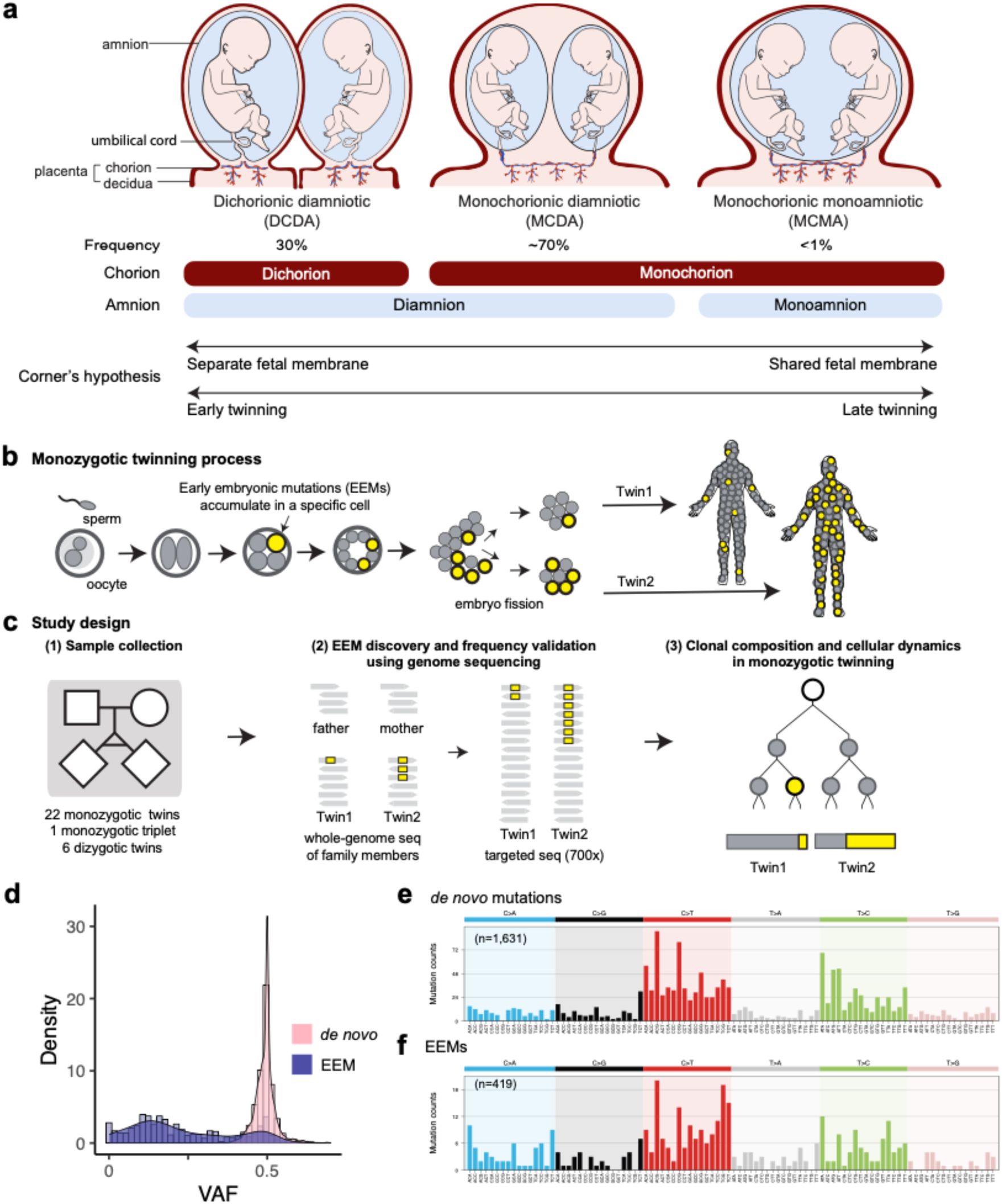
EEMs can be used to infer the clonal dynamics of monozygotic twinning during early embryogenesis. **(a)** Three different fetal membrane types of monozygotic twins. Frequencies of each type are shown, with the numbers of chorions and amnions below. Corner’s hypothesis relating fetal membrane type to the timing of twinning is shown at the bottom. **(b)** EEMs introduced to embryonic cells are stably inherited by daughter cells, irreversibly marking their lineage. Different contributions of embryonic lineages to monozygotic twins are reflected in the varied distribution of EEMs. **(c)** WGS was performed on buccal swab (twins) and peripheral blood (parents). The number of families in this study is shown below the pedigree. Gray rectangles represent sequencing reads, and yellow squares indicate EEMs. EEMs were discovered with WGS and validated by targeted sequencing. VAFs of EEMs were compared between the two twins to infer the clonal dynamics of the twinning process. **(d)** Distribution of VAFs of *de novo* mutations and EEMs detected in monozygotic twins and triplets. **(e,f)** Mutational signatures of **(e)** *de novo* mutations and **(f)** EEMs.

In the 1990s, the pattern of twinning processes was investigated at the cellular level using X-chromosome inactivation as an embryonic lineage marker, which revealed the highest similarities in MCMA twins, followed by MCDA, then DCDA, in overall agreement with Corner’s hypothesis (Chitnis et al. 1999). However, as the X-inactivation starts from the gastrulation (van den Berg et al. 2009), inferring earlier lineages, typically before the twinning event in an embryo, was not possible (Patrat et al. 2020).

Recent genomic studies on somatic mosaicism have revealed that post-zygotic early embryonic mutations (**EEMs**) accumulate from the very earliest embryonic cell divisions of zygotes at a rate of ∼1 mutation per cell per cell division (Park et al. 2021; Coorens et al. 2021a). EEMs are present in a substantial proportion of, but usually not all, cells in postnatal individual somatic tissue. This property allows EEMs to serve as innate molecular barcodes for the reconstruction of early lineages from genome sequencing of postnatal cells or tissues of an individual (**Fig. 1b**)(Park et al. 2021; Ju et al. 2017; Coorens et al. 2021a; Behjati et al. 2014). Recent studies have further reported that EEMs are differentially found between monozygotic twins (Dal et al. 2014; Jonsson et al. 2021; Vadlamudi et al. 2010). By applying systematic strategies to investigate the sharing patterns of EEMs between monozygotic twin pairs, it should be possible to trace early embryonic lineages prior to the twinning process. For example, if an early lineage commonly contributes to both co-twins, the EEMs carried in the ancestral cell (if any) should be shared by the co-twins. Here, we produced whole-genome and deep-targeted sequences from buccal and blood tissues of 22 monozygotic twin families and 1 monozygotic triplet family with the information on chorionicity and amnionicity to test Corner’s hypothesis by retrospectively tracing the clonal dynamics of the human twinning process with EEMs.

## RESULTS

### Whole-genome sequencing for early embryonic mutation profiling in monozygotic twins

To trace the composition of early embryonic lineages in monozygotic twins, we collected buccal and blood tissues from twins and peripheral blood tissues from the parents (**Fig. 1c, Supplementary Table S1**). Our cohort consists of 8 MCMA, 7 MCDA, 7 DCDA monozygotic twins, and 1 dichorionic triamniotic (**DCTA**) monozygotic triplet, as well as 6 DCDA dizygotic twins as controls. Chorions and amnions were verified with prenatal ultrasound by obstetricians as well as histological examination by pathologists. From the twin or triplet siblings, genomic DNA was extracted from buccal swabs, followed by whole-genome sequencing (**WGS**) at 60x read-depth. From the parents, genomic DNA was extracted from peripheral blood and whole-genome sequenced to 30x. Through subtracting DNA variants observed in their parents, we identified non-inherited genomic variants carried in the buccal tissues of the twins, classified into either pre-zygotic *de novo* variants or post-zygotic EEMs (**Methods**, **Supplementary Table S2**). We use the term *de novo* mutations for variants not inherited from either parent, specifically pre-zygotic mutations that were already present in the sperm and oocyte, and EEMs for variants that arise post-zygotically during early embryogenesis.

On average, we detected a median of 67.5 [51-101] pre-zygotic *de novo* variants in monozygotic twins, which is within the range previously reported (Porubsky et al., 2025), ensuring the accuracy of our genome analysis. *De novo* mutations and EEM candidates and their variant allele fractions (**VAFs**) were confirmed by deep-targeted sequencing (700x target coverage). Overall, we identified 1,631 *de novo* mutations and 419 non-redundant EEM events from 22 monozygotic twins and 1 monozygotic triplet. *De novo* mutations had VAF median of 0.50 [range 0.30-0.69] reflecting their full clonal status in a heterozygous diploid genome (**Fig. 1d**). EEM VAF showed a median of 0.12 [range 0-0.64], with a minority of variants presenting as a full clonal mutation in one of the twins, explaining the VAFs closer to 0.5 that are also seen in EEMs (**Fig. 1d**). Mutational spectra of *de novo* mutations and EEMs were decomposed into clock-like signatures (*de novo* mutations: SBS1 14%, and SBS5 86%, cosine similarity = 0.98, **Fig. 1e**; EEMs: SBS1 12%, SBS5 40%, SBS40 39%, and SBS10b 7%, **Fig. 1f**) (Alexandrov et al. 2015), confirming predominantly endogenous mutational processes in the germline and early embryogenesis, as previously reported (Ju et al. 2017; Park et al. 2021; Rahbari et al. 2016).

### Three clonal categories of monozygotic twins

To trace the zygotic origin and early clonal dynamics in each twinning event, non-inherited genomic variants (i.e., *de novo* mutations and EEMs) were deeply compared between co-twins (**Figs. 2a-2d**). In the control dizygotic twins, almost none of the *de novo* variants (0.2%; 2 out of 841) and EEMs (0%; 0 out of 176) were shared between co-twins, reflecting their independent zygotic origins (**Fig. 2a**; **Supplementary Discussion 1, Supplementary Fig. S1**). In contrast, monozygotic twins shared a median of 67.5 *de novo* mutations with a VAF of ∼50% in both twins, consistent with their single zygotic origin (**Figs. 2b-2d**). Furthermore, 45.3% of EEMs were shared in 22 monozygotic twin pairs, suggesting the presence of common downstream lineages that contribute to the somatic tissues of both co-twins.

**Figure 2.**
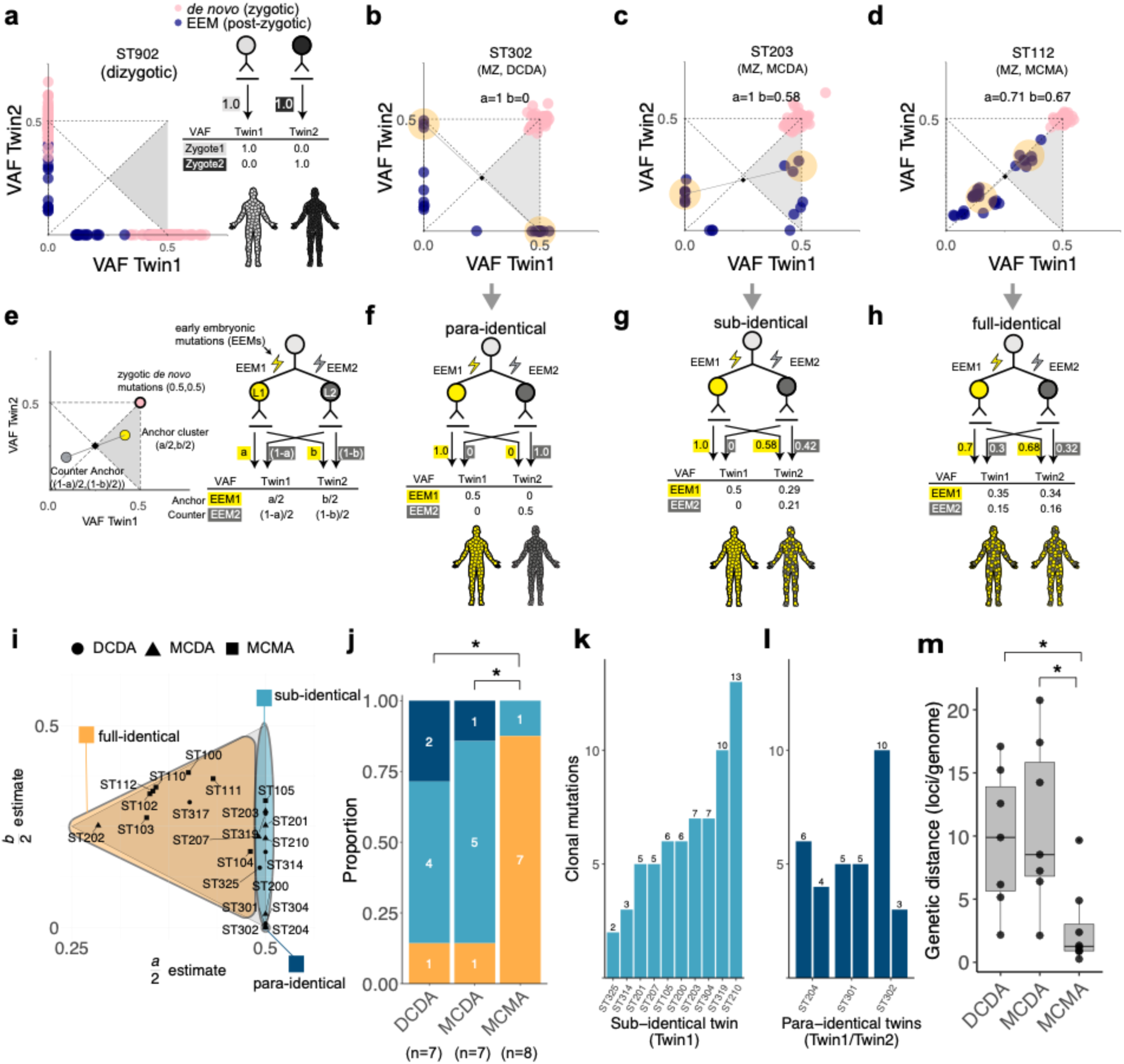
Clonal composition of monozygotic twins. (a) Representative example of dizygotic twins showing mutations and VAFs. Blue dots represent EEMs, and pink dots represent *de novo* mutations. (b-d) Representative examples of monozygotic twins in **(b)** para-identical, **(c)** sub-identical, and **(d)** full-identical categories. Estimated *a* and *b* values are described. Blue dots represent EEMs, pink dots represent *de novo* mutations, and yellow circles indicate anchor and counter-anchor mutation clusters. **(e)** Mathematical framework for clonal composition of monozygotic twins. The anchor (yellow circle) and counter-anchor (gray circle) clusters, representing the two earliest detectable embryonic lineages (L1 and L2), are shown with their VAFs in a twin pair. The black diamond represents the expected midpoint of anchor and counter-anchor clusters at (0.25, 0.25), and the pink circle indicates the expected position of *de novo* mutations at (0.5,0.5). The contributions of L1 to the somatic tissues of Twin 1 and Twin 2 are defined as *a* and *b*, respectively. The contribution of the earliest lineages to twins and expected VAFs of corresponding mutations are summarized with respect to *a* and *b*. Twin 1 is defined as the twin with the higher L1 contribution, with anchor mutations consistently located in the gray triangle. **(f-h)** Contribution of the two earliest cells in three clonal categories of monozygotic twins: **(f)** para-identical, **(g)** sub-identical, **(h)** full-identical, each corresponding to the twins described in panels (b), (c), and (d), respectively. **(i)** Positions of anchor cluster (a/2, b/2) 22 monozygotic twins. DCDA, MCDA, and MCMA twins are shown as circles, triangles, and squares, respectively. **(j)** Proportion of clonal categories in 22 monozygotic twins with different fetal membrane configurations. **(k)** The number of clonal mutations unique to Twin 1 of 10 sub-identical twins, with counts shown above each bar. **(l)** The number of clonal mutations in 3 para-identical twin pairs, with counts shown above each bar. (m) Distribution of genetic distances between 22 monozygotic twins across different fetal membrane subtypes. \**P* < 0.05.

To systematically evaluate the clonal relationship in monozygotic co-twins based on early lineages, we constructed a cell genealogy model focusing on the two earliest informative ancestral cells (referred to as **L1** and **L2** throughout the manuscript) contributing to the somatic tissues of each twin in defined ratios (**a** and **b** for the contributions of L1 to Twin 1 and Twin 2, respectively; **Fig. 2e**; **Supplementary Fig. S2, Methods**). This model is scrutinized by a specific cluster of EEMs, termed ‘**anchor cluster**’, which exhibit dominant VAFs across the two monozygotic co-twins in aggregate (i.e., ≥50% when summing VAFs from both twins, falling within the gray triangle of the plot; **Fig. 2e**). In principle, if each embryonic cell acquires its unique detectable post-zygotic mutation, the mutations in the anchor cluster correspond to the EEMs carried by the common ancestor of the dominant lineages (**L1**), the ancestral cell that gave rise to the majority, but not all, of the cells comprising the embryonic proper (or soma) of twin embryos as a whole. At the same time, our model predicts the existence of a ‘**counter-anchor cluster**’, which appears as point-symmetric counterparts relative to the central point (0.25,0.25) in the two-dimensional plot (**Fig. 2e**). These mutations represent those acquired in the common ancestor of the minor lineages (**L2**), since the combined contribution of L1 and L2 (equivalent to twice the VAF of the corresponding mutations) should be 100% in each twin. For consistency, we define Twin 1 as showing an anchor cluster with higher VAFs (**a** ≥ **b**; **Fig. 2e**).

Of note, since not all embryonic cells carry their own detectable mutations (as not all cell divisions lead to detectable EEM accumulation), anchor and/or counter-anchor EEMs may be absent in real-world datasets. In the genome sequences of 22 monozygotic twin families, 17 (77%) and 22 (100%) exhibited anchor and counter-anchor EEM clusters, respectively (**Supplementary Table S3**), suggesting that our EEM-based lineage tracing is possible in monozygotic twins (Ju et al. 2017; Park et al. 2021).

The VAFs of EEMs in anchor clusters revealed three distinct clonal lineage categories among monozygotic twins. First, in twins defined as **para-identical**, somatic cells of Twin 1 and Twin 2 originated exclusively from L1 and L2, respectively (a = 1, b = 0; **Figs. 2b, 2f**), indicating two distinct contralateral founder cells in early embryonic lineages. Second, in twins classified as **sub-identical**, Twin 1 was derived exclusively from L1, whereas Twin 2 arose from both L1 and L2 (a=1, b<1; **Figs. 2c, 2g**). The pattern suggests that the entire embryonic lineage for Twin 1 is nested within that of Twin 2, directly implying that the founding cell of Twin 1 emerged several cell cycles later than that of Twin 2. Lastly, in those classified as **full-identical**, the contributions of L1 (and L2) were largely comparable between Twin 1 and Twin 2 (a ∼= b; **Figs. 2d, 2h**), suggesting that the physical separation of the twins occurred at a later stage, after the establishment of multiple early embryonic cells (lineages) at the equilibrium within the common embryo.

### Clonal categories associated with the amnionicity of monozygotic twins

Collectively, 22 monozygotic twins in our cohort comprised 3 para-identical, 10 sub-identical, and 9 full-identical twins based on their anchor cluster analysis (**Fig. 2i, Supplementary Fig. S3**). To associate with Corner’s hypothesis, we then correlated the clonal categories of monozygotic twins with fetal membrane subtypes and revealed a relationship between fetal membranes and clonal categories. Twin subtype distributions differed significantly across groups (**Fig. 2j**; MCMA, MCDA, DCDA; 3×3 Fisher Monte Carlo simulation, *P* = 0.018). MCMA twins were almost exclusively full-identical (87.5%), with only a single sub-identical pair and no para-identical pairs. By contrast, MCDA and DCDA twins showed nearly identical distributions, both dominated by sub-identical and para-identical categories, with only 14.3% full-identical each. Pairwise comparisons confirmed this pattern (**Fig. 2j**). MCMA was significantly different from both MCDA (Fisher’s exact test, Bonferroni correction, *P* = 0.036) and DCDA (*P* = 0.045), whereas MCDA and DCDA did not differ (*P* = 1). Thus, when comparing across chorionicity, monochorionic twins as a group (MCMA+MCDA) appeared different from dichorionic twins (DCDA; *P* = 0.017), but this effect was entirely attributable to the MCMA subgroup. When directly comparing by amnionicity, the distinction was even clearer with monoamnionic twins (MCMA) with majority of twin pairs with full-identical (87.5%), whereas diamniotic twins (MCDA+DCDA) were rarely full-identical (14%, *P* = 0.003). In addition, para-identical twins with two independent ancestral cells were observed exclusively among diamniotic twins, although the association did not reach statistical significance, likely due to the limited cohort size (0% vs. 21.4%, *P* = 0.26). Our results collectively suggest that the clonality of twins is tightly associated with amnionicity but not chorionicity.

We hypothesize that clonal categories are closely linked to the timing of physical twinning. For example, if an embryo’s effective cell population is minimal (i.e., 2-cell stage), then complete segregation of the lineages (e.g., L1 or L2) leading to para-identical twins is inevitable. In contrast, once the embryo reaches a large effective cell population size, complete segregation of upstream lineage origins becomes highly improbable, leading to full-identical twins. Between the two extremes, if twinning occurs at an intermediate stage, one twin originates entirely from a single upstream lineage while the other inherits a mixture of lineages, resulting in sub-identical twins. Overall, this implies that the physical separation giving rise to para-identical twins must occur earlier than in other clonal categories. In addition, the presence of shared anchor cluster EEMs, observed in full-identical twins, would not be compatible with twinning at the earliest embryonic stage (i.e., the 2-cell stage), but rather at a later developmental stage than that inferred for the other clonal categories. Corner’s hypothesis proposed that both fetal membranes (chorions and amnions) are correlated with the timing of twinning. Our results refine this model by showing that only the amnion, but not the chorion, is tightly correlated with the clonality of twins. As amnions are formed around day 8-12 in human embryos (Oldak et al. 2023), our findings suggest that the timing of full-identical twinning would occur after day 8-12.

Among the three clonal categories of monozygotic twins, sub-identical twins clearly show a temporal order of founder cells between Twin 1 and Twin 2. Because endogenous somatic mutations acquired during embryogenesis accumulate in a clock-like manner (approximately 1.2 mutations per cell per cell division (Park et al. 2021)), the number of clonal mutations unique to Twin 1 reflects the number of cell divisions separating the founder cells of Twin 1 and Twin 2. From the 10 sub-identical twin pairs, we observed a median of 6 clonal mutation differences between Twin 1 and Twin 2 (range 2–13; **Fig. 2k**), suggesting that the founder of Twin 1 emerged approximately 5 cell divisions after that of Twin 2. Furthermore, in 3 para-identical twin pairs, the founder cells of Twin 1 and Twin 2 carried a median of 5 unique mutations, suggesting that the founder cells were at least 4 cell generations downstream from the common ancestral cell, often the fertilized egg (**Fig. 2l**).

With recent studies showing that monozygotic twins are not genetically identical (Jonsson et al. 2021), we calculated the genetic distance between twins using EEMs and compared them across fetal membrane subtypes (**Fig. 2m**). The mean genetic distances were 2.69, 10.96, and 9.74 for MCMA, MCDA, and DCDA twins, respectively, with statistically significant differences observed among the groups (Kruskal-Wallis test, *H*(2) = 9.49, *P* = 0.0087). The post-hoc subgroup analysis again showed significance only between monoamniotic and diamniotic twins (MCMA vs MCDA, *P* = 0.0058; MCMA vs DCDA, *P* = 0.0123), with no significant difference within diamniotic twins (DCDA vs MCDA, *P* = 0.81).

### Shared clonal compositions in cord blood of monochorionic twins

From 14 monozygotic twin families, we collected cord blood samples at birth in addition to buccal tissues. Genomic DNA from cord blood was also whole-genome sequenced to 60x. VAFs of EEMs were generally consistent between buccal and cord blood in all available dichorionic twins (n=3; **Fig. 3a**), with clonal categories inferred from cord blood matching those from buccal tissues (**Fig. 3b, Supplementary Fig. S4**). In contrast, in monochorionic twins (n=11), VAFs of EEMs substantially differed from those in buccal tissues (**Fig. 3c**), but were concordant between the cord bloods of Twin 1 and Twin 2 (**Fig. 3d**), suggesting extensive blood exchange *in utero*. Consequently, clonal categories estimated from cord blood were uninformative, showing full-identical-like patterns in all monochorionic twins (**Fig. 3d, Supplementary Fig. S2**), regardless of the categories inferred from buccal tissues. These findings align with the shared circulatory system of monochorionic placentas.

**Figure 3.**
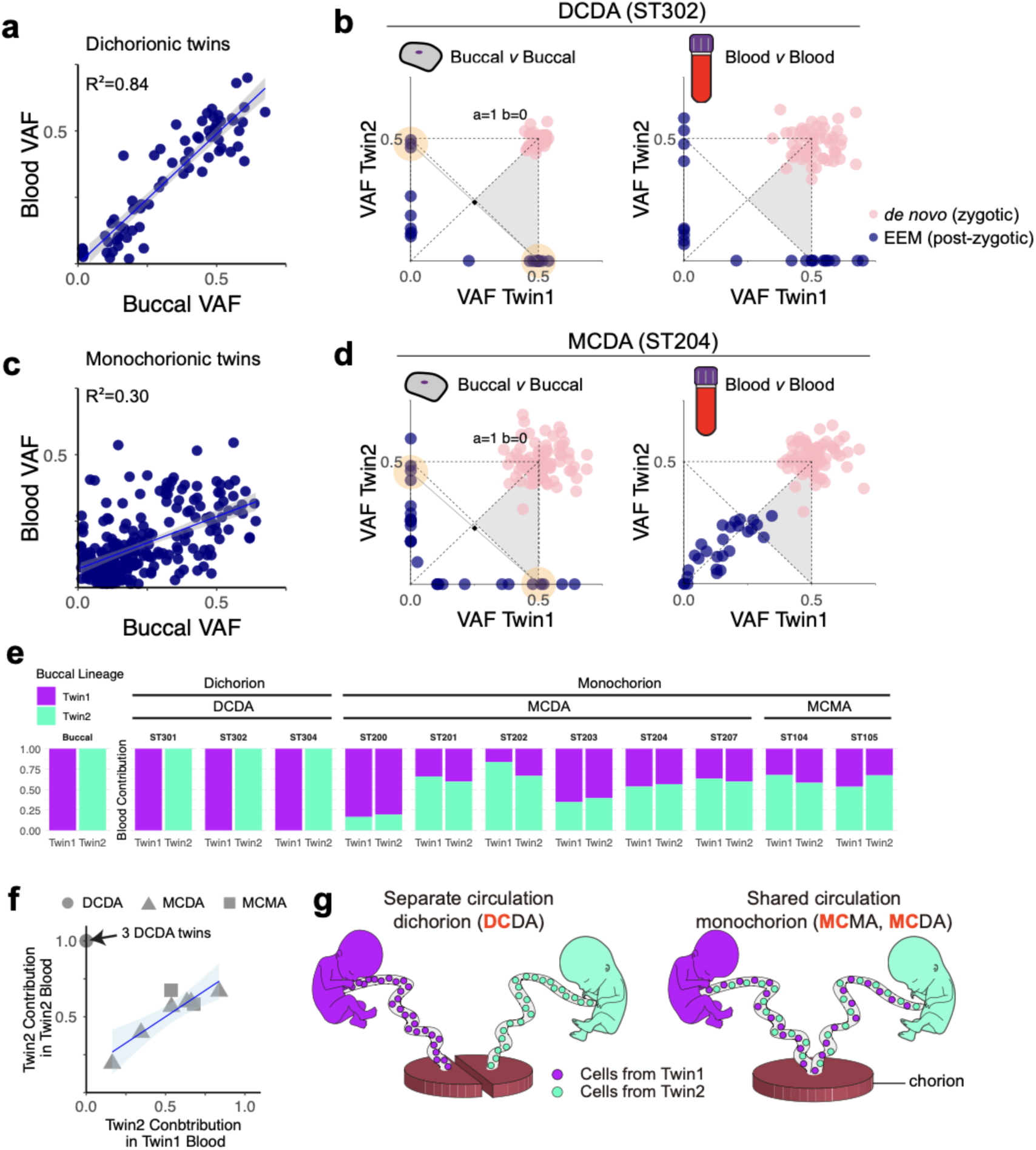
Hematopoietic cell mixing in monochorionic twins. **(a)** Correlation between blood and buccal VAFs in three DCDA twins. Blue dots represent EEMs. The blue line shows the linear regression (y = 0.98x + 0.0, R² = 0.84), and the shaded area represents the 95% confidence interval. **(b)** Comparison of VAFs of EEMs between buccal (left) and blood (right) samples in a DCDA twin pair. Blue dots represent EEMs, and pink dots represent *de novo* mutations. **(c)** Correlation between blood and buccal VAFs in 11 monochorionic (MCDA and MCMA) twins. Blue dots represent EEMs. The blue line shows linear regression (y = 0.39x + 0.07, R² = 0.30), and the shaded area represents the 95% confidence interval. **(d)** Comparison of VAFs of EEMs between buccal (left) and blood (right) samples in an MCDA twin pair. Blue dots represent EEMs, and pink dots represent *de novo* mutations. **(e)** Blood mixing ratios estimated by maximum likelihood in 11 monozygotic twin pairs. Fetal membrane types are shown on the top. **(f)** Contribution of Twin 2 to the blood of Twin 1 and Twin 2. The blue line shows the linear regression of MCMA and MCDA twin data (y=0.69x + 0.15, R² = 0.80) with the shaded area indicating the 95% confidence interval. The black arrow indicates three DCDA twin data. DCDA, MCDA, and MCMA twins are shown as circles, triangles, and squares, respectively. **(g)** Illustration showing unexchanged hematopoietic systems in dichorionic twins (DCDA; left) and hematopoietic cells exchanged through the common placenta in monochorionic twins (MCMA and MCDA; right).

Of the 14 monozygotic twin pairs with both buccal and blood DNA, 11 twin pairs had at least one unique EEM restricted to one twin (either sub-identical or para-identical) that could be informative for quantifying the relative contributions of Twin 1 and Twin 2 to the pool of cord bloods. All monochorionic twins (MCMA and MCDA) showed varying degrees of hematopoietic cell mixing, with the major contributor accounting for 56%-83%, often deviating from an equal 50% contribution from both twins (**Fig. 3e**). Here, blood cells from the two twins showed approximately similar degrees of contribution from one of the twins (**Fig. 3f**), also supporting the idea that the two twins have a common hematopoietic system that is well mixed. In contrast, as expected, DCDA twins showed no mixing of hematopoietic cells between the twins (**Fig. 3f**). Our findings demonstrate that monochorionic monozygotic twins share a common clonal composition in their cord blood, independent of the early embryonic clonal dynamics that shape solid tissue formation (**Fig. 3g**).

### Confirming clonal categories using single-cell analysis

In this study, we primarily analyzed DNA extracted from buccal swabs (bulk tissues) to infer clonal categories, as access to other normal tissues, apart from cord blood, was limited due to ethical considerations. However, we recognized that single-cell-resolution lineage tracing is feasible in dichorionic twins (DCDA) using cord blood, as these tissues permit single-cell isolation and largely preserve the clonal categories reconstructed from buccal tissues (**Figs. 3a, 3b**). To confirm the clonal categories at the single-cell level, peripheral blood mononuclear cells (**PBMCs**) were isolated from two DCDA twins (ST317 (full-identical) and ST314 (sub-identical)) via fluorescence-activated cell sorting (**FACS**) technique. The whole genomes of 182 single PBMCs were amplified using the multiple displacement amplification (**MDA**) method (Dean et al. 2002), followed by deep-targeted sequencing for EEMs and *de novo* mutations discovered in buccal WGS (**Fig. 4a**). Accounting for allelic or locus dropouts inherent to single-genome amplification (estimated to be 23% and 46% in our dataset, respectively; (Woodworth et al. 2017)), we reconstructed single-cell-resolution developmental phylogenies in PBMCs, which validated the clonal categories inferred from the whole-genome sequences of the bulk tissues.

**Figure 4.**
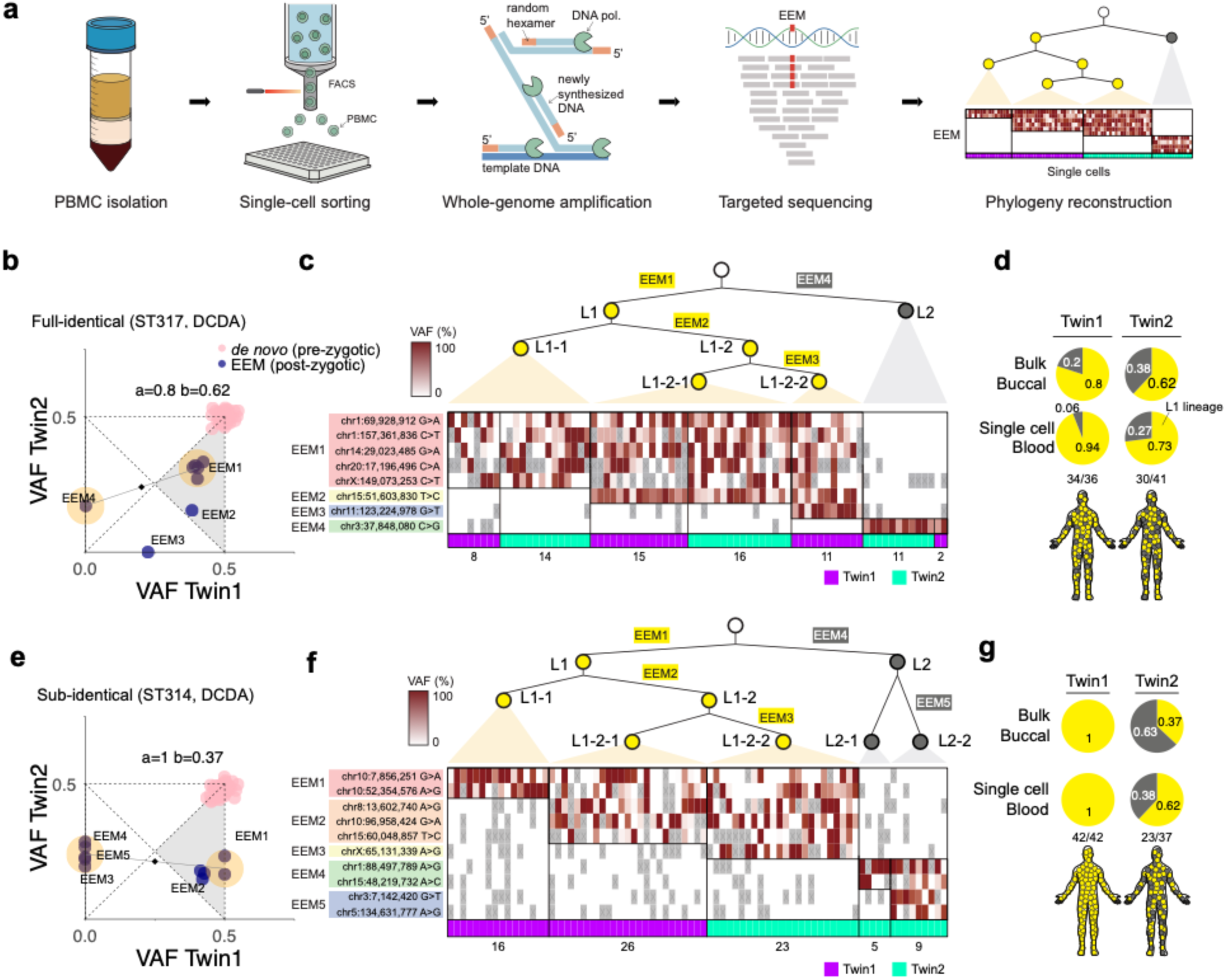
Single-cell lineage tracing using cord blood of dichorionic monozygotic twins. **(a)** Experimental design for single-cell lineage tracing in dichorionic monozygotic twins. Phylogenies were reconstructed based on EEMs identified in single PBMCs from each twin, which underwent FACS sorting, whole-genome amplification, and targeted sequencing. **(b)** VAFs of EEMs and *de novo* mutations detected in bulk buccal tissues of ST317 twins (full-identical, DCDA). Blue dots represent EEMs, and pink dots represent *de novo* mutations. Yellow circles indicate anchor and counter-anchor mutation clusters. **(c)** Reconstruction of developmental lineages based on EEM VAFs in single PBMCs from ST317. The heatmap is colored according to EEM VAFs in a red gradient, while ray boxes marked with ‘x’ indicate absence of informative reads due to dropouts. Single cells from Twin 1 and Twin 2 are marked with purple and cyan, respectively. The number of cells in each group is indicated below the boxes. **(d)** Estimated contribution of L1 lineages to bulk buccal tissue and PBMCs in ST317 twins. **(e)** VAFs of EEMs and *de novo* mutations detected in buccal tissues of ST314 twins (sub-identical, DCDA). **(f)** Reconstruction of developmental lineages based on EEM VAFs in single PBMCs from ST314. **(g)** Estimated contribution of L1 lineages to bulk buccal tissue and PBMCs in ST314 twins.

In ST317, a full-identical DCDA twin, the phylogeny reconstructed from single PBMCs was completely concordant with the clonal histories derived from buccal WGS (**Figs. 4b-4d**). The 5 anchor EEMs (EEM1) and 1 counter-anchor EEM (EEM4), inferred to be carried by the common ancestors of L1 and L2, respectively, were mutually exclusive in 36 PBMCs from Twin 1 and 41 PBMCs from Twin 2 (**Figs. 4b, 4c**). Furthermore, both L1 and L2 lineages were detected in the PBMCs of both twins, with contributions comparable to those in buccal tissues, confirming their full-identical status (**Figs. 4c, 4d**). In ST314, a sub-identical DCDA twin, single PBMC sequencing further resolved early embryonic lineages while validating the clonal categories defined from buccal WGS. The buccal WGS identified 2 anchor EEMs (EEM1) and 5 counter-anchor EEMs (**Fig. 4e**). However, in 42 and 37 PBMCs from Twin 1 and Twin 2, respectively, EEM1 was only mutually exclusive with 4 counter-anchor EEMs (EEM4 and EEM5) but co-occurred with 1 counter-anchor EEM (EEM3; chrX:65,131,339, A>G; **Fig. 4f**), suggesting that this mutation occurred in a downstream lineage of L1 but coincidentally overlapped with the counter-anchor cluster in bulk WGS. Aside from this additional insight, developmental phylogenies of 79 PBMCs clearly confirmed the sub-identical pattern predicted from bulk buccal tissues (**Figs. 4f, 4g**). Collectively, our findings validate the bulk-tissue-based inference of clonal categories defined by the two earliest ancestral cells (i.e., L1 and L2), despite insensitivities for decomposing more downstream lineages of L1 and L2.

### Two consecutive twinning processes for monozygotic triplets

The WGS of a DCTA monozygotic triplet family (ST501) identified 70 *de novo* mutations and 33 EEMs across the triplets. Interestingly, clonal relationships in early development, inferred from the sharing pattern of EEMs, were not equal among the three triplets (**Fig. 5a**). Using our cell genealogy model, we identified sub-identical twin relationships in all three possible pairwise combinations (**Fig. 5b**). Furthermore, Triplet 1 and Triplet 2 shared a later most recent common ancestor (**MRCA**) cell than the other pairs (**Fig. 5c**). Our findings showed that both Triplets 1 and 2 were subclonal to Triplet 3, placing Triplets 1, 2 combined were branched out first from Triplet in a subclonal fashion. Then, Triplet 1 was separated from Triplet 2 by a secondary sub-identical twinning process (**Fig. 5c**). Of note, given the abovementioned association between clonal categories and amnionicity (**Fig. 2j**), the two consecutive sub-identical twinning processes were compatible with three separate amnions in the triplet (**Figs. 5a, 5c**).

**Figure 5.**
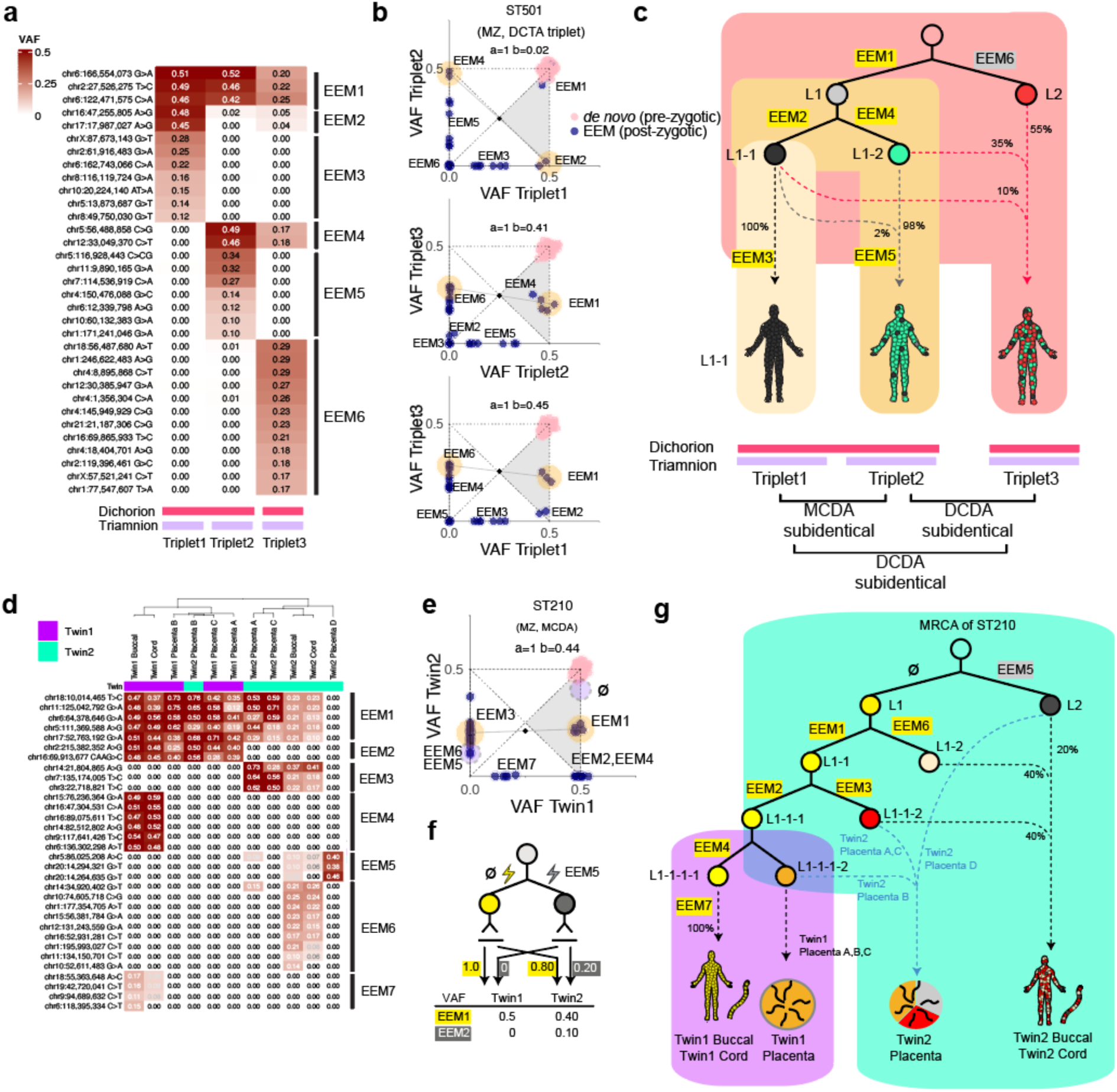
Clonal composition of a monozygotic triplet family and extraembryonic lineage of a monozygotic twin family. **(a)** A heatmap of VAFs of EEMs of the monozygotic triplets. VAFs are represented as gradients in red. Mutation clusters are shown on the right side of the heatmap corresponding to each cluster on the x-y coordinates shown in (b). The fetal membrane configuration of the triplets is shown under the heatmap. **(b)** Two-way VAF comparisons between the monozygotic triplets. Anchor and counter-anchor clusters are shown with a yellow background. Each mutation cluster from the heatmap in (a) is shown for each x-y plot. **(c)** Estimated lineage tree and contribution to each triplet with respective fractions. The chorion and amnion relationship between the triplets is drawn below each triplet. MRCA: most recent common ancestor **(d)** VAF heatmap of multiple tissues (buccal epithelial cells, umbilical cord, placenta) of ST210 (MCDA sub-identical twin). Twin of origin is shown in purple for Twin 1 and cyan for Twin 2. EEM clusters are grouped on the right side of the plot. **(e)** EEMs shown on the x-y plot result from buccal whole-genome sequencing of ST210. Yellow circles in the 2d plot represent the anchor and counter-anchor mutation clusters estimated from buccal WGS. The blue dotted area represents the new anchor and counter-anchor mutations proposed based on the VAF heatmap in (d). ø represents no mutation detected in the specific cell division lineage. **(f)** Updated lineage model with new anchor and counter anchor mutations analysis. **(g)** Lineage history of placenta, umbilical cord, and buccal epithelial cells of ST210.

### Twin lineage separation can precede embryonic and extra-embryonic fate decisions

Finally, in a sub-identical MCDA twin (ST210), we were able to extend the early embryonic lineages inferred from buccal epithelial cells to extraembryonic tissues, including the umbilical cord and placenta, owing to the availability of these samples. From placental tissues, we performed laser-capture microdissection (**LCM**) of four micropatches from each twin, each measuring <1 mm² and containing <3,000 cells. The bulk umbilical cord samples and placental micro-patches were then whole-genome sequenced to ∼24x, followed by confirmation of EEMs identified from bulk buccal epithelial cells.

Consistent with our single-cell genome observations (**Figs. 3b–3g**), clustering of EEMs from additional tissues, including two umbilical cords and seven LCM-derived placental samples, resolved early clonal dynamics with greater temporal resolution than buccal tissues alone, while maintaining a consistent clonal category between the co-twins (**Figs. 5d, 5f**). Here, 15 EEMs previously clustered within or adjacent to the counter-anchor group were further resolved, with EEM5 identified as the true counter-anchor cluster and an unidentified anchor cluster, further refining our lineage model (**Fig. 5e**). The deeper phylogeny revealed that the early clonal composition of umbilical cords is nearly identical to that of buccal epithelial tissues, suggesting that the umbilical cord and the embryo proper diverged at a relatively late developmental stage (**Figs. 5d, 5g**). However, the origin of placental tissues was slightly different. In Twin 2, the clonal origin of four LCM patches was substantially different from each other (Placenta A and C from L1-1-2, Placenta B from L1-1-1-2, and Placenta D from L2 predominantly), suggesting clonal expansion for a restricted region (LCM patch), consistent with a previous observation (**Figs. 5d, 5g**)(Coorens et al. 2021b). In addition, the subclonal divergence between Twin 1 and Twin 2 occurred prior to the complete cell fate segregation between extraembryonic and embryonic lineages (**Fig. 5g**), as L1-1-1 still contributed to both tissue types. Similarly, the clonal distance from buccal to placental tissues of Twin 1 was smaller than the distance to buccal or placental tissues of Twin 2. These observations suggest that the sub-identical twinning event in this MCDA twin occurred prior to embryonic day 5–6, before the segregation of trophoblasts (progenitors of the chorion) from embryoblasts (progenitors of the embryo proper). Notably, the presence of monochorionicity despite multiple clonal origins (at least from L2, L1-1-2, L1-1-1-2) supports our earlier conclusion that chorionicity is not necessarily associated with clonal history.

## DISCUSSION

The timing of monozygotic twinning and its relationship to fetal membranes has remained an unresolved question since Corner’s hypothesis. Here, we systematically analyzed the clonal architecture of monozygotic twins with defined membrane subtypes to estimate the timing of twinning events using whole-genome sequencing. By comparing VAFs of EEMs between co-twins, we revealed three clonal categories, para-identical, sub-identical, and full-identical twins, based on anchor and counter-anchor clusters. Our data show that amnionicity, but not chorionicity, is closely linked to clonal categories. These findings refine and extend Corner’s hypothesis, providing direct evidence from clonal dynamics and offering a new perspective to the conventional anatomical classification of DCDA, MCDA, and MCMA based on the fetal membranes of monozygotic twins. Technically, by combining clonality from solid and blood tissues, we provide a new framework for understanding monozygotic twinning in relation to their fetal membranes (**Fig. 6**). Notably, our estimation of twinning is based on the lineage separation of the ancestral cells of the co-twins, which may not precisely coincide with their physical separation. Nevertheless, given the challenging nature of studying the human embryo, our study provides the most detailed lineage reconstruction of twins to date, offering the strongest evidence yet for inferring the timing of monozygotic twinning.

**Figure 6.**
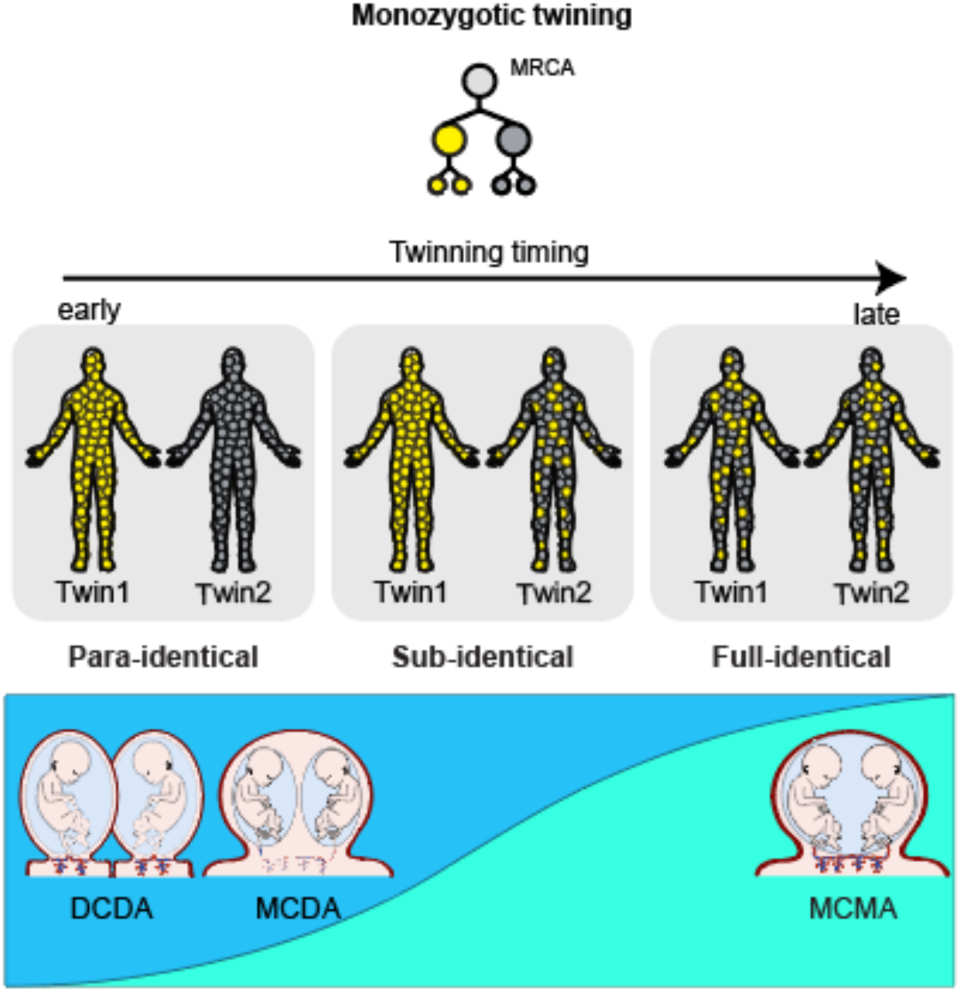
An updated twinning model. Clonal relationships in monozygotic twins are shown in relation to the timing of twinning. Colors represent specific cell lineages. Three possible clonal relationships in the monozygotic twins (para-identical, sub-identical, and full-identical) are represented. Frequencies of each clonal type and amnionicity are shown below.

Conjoined amnion (monoamnion) was predominantly observed in full-identical twins, in which the clonal compositions of the two individuals were highly similar. In contrast, the clonal basis of conjoined chorions (mono chorions) remains unresolved. Addressing this question will require analysis of clonal dynamics in both somatic and extraembryonic tissues, yet in the present study placental material was available from only a single twin pair. Large-scale prospective studies will therefore be essential, as placental tissue collection is rarely incorporated into routine clinical practice.

Monochorionic twins always exhibited shared circulation, with one co-twin predominantly contributing white blood cells to the pool of cord bloods to the extent of 56%-83% at birth (**Fig. 3e**). In principle, two non-exclusive mechanisms may underlie the observed blood cell asymmetry: (1) asymmetric blood flow between monozygotic twins *in utero* that persists until birth, often leading to twin-to-twin transfusion syndrome (**TTTS**)(Lewi et al. 2008); and (2) inter-twin exchange of hematopoietic stem cells during embryonic development, which subsequently engraft in the bone marrow, the principal site of fetal hematopoiesis after the 20^th^ week of gestation, beyond the yolk sac, liver, and spleen (De La Garza et al. 2017). In our previous observation of monochorionic dizygotic twins, the blood cell asymmetry resulting from a monochorionic blood exchange *in utero* remained stable for a long time, observed in at least 5-year-old co-twins (Yoon et al. 2024), supporting the latter scenario. Notably, despite the asymmetric contribution of cord blood circulation, none of our monochorionic twins’ pregnancies were complicated by TTTS. Studies have shown that monozygotic twins can develop hematological malignancies with identical EEM, which is shared through the transplacental transmission of hematopoietic stem cells, demonstrating the consequences of shared circulation between monochorionic monozygotic twins (Sousos et al. 2022; Hong et al. 2008).

Although bulk buccal tissue comparisons, the primary approach in this study, were informative for assessing the relative contributions of the earliest two cells in twins, single-cell phylogenies provided finer resolution of clonal dynamics. Incorporating placental tissues further extended these insights to extraembryonic lineages, a critical component of embryogenesis. Future large-scale genomic studies of monozygotic twins, integrating single-cell data from both embryonic and extraembryonic tissues, will be essential to resolve cellular dynamics in twinning and to refine clonal categories in human monozygotic twin development.

## Acknowledgements

This work was supported by the National Institutes of Health (F30 HD106744 to C.J.Y.), the National Research Foundation Korea (NRF-2020R1A3B2078973 to Y.S.J.), the Korea Health Technology R&D Project through Korea Health Industry Development Institute, which was funded by the Ministry of Health & Welfare of Korea (RS-2024-00440005 to Y.S.J.), and the Suh Kyungbae Foundation (SUHF-18010082 to Y.S.J.).

## Author contributions

Conceptualization: C.J.Y., J.K.J. and Y.S.J. Methodology and formal analysis: C.J.Y. and C.H.N., Mutational signature analysis: J.L., Resources: E.S.C., J.H.B., H.K., Y.M.J., S.M.L., C.D., E.M., S.W., R.K., J.S., J.K.J. Computational Resources: J.L., J.W.P., Writing: C.J.Y., C.H.N., and Y.S.J., Supervision: O.L.G., M.G., J.K.J., and Y.S.J. All authors have reviewed and approved the manuscript.

