## Supplementary material for "Clonal dynamics of monozygotic twinning in early human embryogenesis": Material Methods, Supplementary Figures

**Supplementary Discussion 1: Lineage history of dizygotic twins**

One dizygotic twin pair (ST900) shows two *de novo* mutations that are shared between the two dizygotic twins (**Supplementary Fig. S1a**). Dizygotic twins are derived from two separate zygotes thus, *de novo* mutations that are acquired from the sperm lineage or oocyte lineage in the parents are usually not shared if the two sperms and two oocytes forming each twin are derived from distinct lineages. The presence of the two shared *de novo* mutations indicates a shared lineage in either the sperm and/or oocyte lineages in the two dizygotic twins (**Supplementary Fig. S1b**). When we phased these two shared *de novo* mutations to the nearest parental SNPs, they were phased to a maternally inherited SNP, indicating their maternal origin. Thus, we concluded that the two shared *de novo* mutations were acquired in the shared oocyte lineages of the two dizygotic twins. Since each oocyte receives half of the genetic material (n) from its precursor (2n), only ¼ of the entire maternal (2n) genome is shared between any two oocytes. Thus, we inferred that at the most common recent ancestor of the two oocytes, there were 8 mutations that were acquired before the two oocytes' lineage tree diverged. As shown in prior studies on genetic siblings from multi-sibling families (Jónsson et al. 2018) and our previous study on monochorionic dizygotic twins (Yoon et al. 2024), these results demonstrate that *de novo* mutations shared between genetic siblings of dizygotic twins are common. Further studies will be needed to quantify the rate and how these differ between twins conceived naturally and with assisted reproductive technologies.

### Materials and Methods

**Sample collection, whole-genome sequencing, and quality control**

Twin families (22 monozygotic twins, 1 monozygotic triplet, and 6 dizygotic twins) were recruited from the Department of Obstetrics and Gynecology at Seoul National University Hospital under the IRB protocol (SNUH IRB H-1311-045-533, KAIST IRB KH2019-134). Fetal membranes were confirmed with prenatal ultrasound examinations and microscopic examination of amnions and chorions after birth by pathologists. Written consent was obtained from the parents at the time of sample collection. Peripheral blood was obtained from the parents and the twins. Buccal epithelial cells were obtained from the twins using the AccuBuccal Collection Kit (Accugene). DNA was extracted using the DNeasy Blood and Tissue kit (Qiagen). Truseq PCR-free library (Illumina) was generated for the blood DNA (>1μg). Truseq Nano library (Illumina) was generated for the buccal epithelial DNA (< 1μg). Whole genome sequencing was conducted at 30x for the parents' blood DNA and 60x for the twins' buccal DNA on an Illumina NovaSeq

6000. Raw FASTQ files were aligned to the human reference genome GRCh38 with bwa mem (Li 2013) and deduplicated with samblaster (Faust and Hall 2014). Indels were realigned with Genome Analysis Tool Kit (GATK) v3.8 IndelRealigner (McKenna et al. 2010). TrioMix v0.0.2a (Yoon et al. 2022) was used to estimate contamination within family members. VerifyBamID2 v2.0.1(Zhang et al. 2020) was used to estimate contamination from unrelated individuals. Twin Buccal DNA with estimated contamination < 1% were used for further analysis.

**EEM detection**

SNPs and indels were called by using Varscan2 (v.2.4.4) (Koboldt et al. 2012) with family member's bam file as trios as inputs. Each twin's variants were called independently as two separate trios and then merged. Non-inherited variants (*de novo* mutations and early embryonic mutations (EEMs)) were obtained by removing variants that had any variant read in the parents. Variants were required to have at least 10 read depth in the father, the mother, and the twins. Reads in poorly mapped regions were filtered by removing loci where total reads with soft or hard clips were greater than 20 % or regions with surrounding ± 150bp with greater than 5% soft or clipped read fractions. Also, loci with more than 10 reads with a mapping quality of 0 and a median mapping quality of less than 40 were removed. Variants were also removed if the difference in mapping quality between the variant read and reference read was greater than 20, as these likely resulted from poor read mapping. Variants with a median NM tag value difference between variant read and reference reads greater than 5 were removed. Variant Effect Predictor (VEP v.93) was used to annotate variants with population allele frequencies from gnomAD v.2.0.1 (McLaren et al. 2016; Karczewski et al. 2020). Variants present in greater than 1% in any population group (max AF) were removed. Additionally, panels of normal bam files from all parents included in this study and all other twin pairs were used to filter out artifacts from common SNPs. We used depth from these heterozygous loci in the twins (where one of the parents is homozygous reference and the other parent is homozygous alternative genotype) to estimate the depth and VAF distribution of each child's WGS. Variants falling outside of 0.5-99.5 percentiles of sequencing depth distributions were removed to avoid copy number variation regions. Variants where the combined VAF of the two twins was greater than 0.1 were used for final target sequencing bait design for validation and further analysis.

**Target sequencing of twin families**

Integrative DNA Technologies (IDT) Discovery Pool custom baits for each twin family were designed for the putative EEM and de novo mutation sites discovered with whole-genome sequencing. Custom probes were hybridized to the DNA libraries and were sequenced with Illumina NovaSeq 6000 with approximately ~700x coverage.

**Estimation of overdispersion in whole-genome sequencing and targeted sequencing**

We selected SNVs where one of the parents is a homozygous reference genotype and the other parent is a heterozygous genotype in the autosome. The children's genotype must be heterozygous according to Mendel's law. For these loci where the 'true' VAFs are known at 0.5, we fitted our whole-genome sequencing and targeted sequencing results to estimate the overdispersion factors in the beta binomial distribution, resulting in $\rho_{WGS}$= 0.0002568, $\rho_{TGS}$= 0.0008809.

**Variant curation**

In the final variant curation, reads from both whole-genome sequencing and target sequencing were compared. If the target sequencing depth was lower than whole-genome sequencing in at least one of the twins, whole-genome sequencing data was used for that locus. If a variant has a nearby phasable SNP and the total number of reads containing the variant and the SNP with target-sequencing is greater than whole-genome sequencing depth, adjusted VAF using only the reads containing the variant and the SNP were used for VAF calculation. If the EEMs were completely phased to the nearby SNP (i.e., only two haplotypes exist), the EEM's VAFs are adjusted to reflect the VAF of clonal mutations, 0.5. For male children, VAFs of mutations occurring outside of the pseudoautosomal regions (PARs) of the X chromosome were adjusted as males are haploid. All *de novo* mutations and EEMs were manually reviewed with the Integrative Genomics Viewer (IGV).

Clonality of a mutation was determined using a two-tailed confidence interval of 99.9 percentile with the expectation of a heterozygous variant with 50% allele frequency with a beta binomial distribution and VAF > 0.3. Dispersion parameters $\rho_{WGS}$= 0.0002568 and $\rho_{TGS}$= 0.0008809 were used. For monozygotic twins, a mutation is determined to be *de novo* if the mutation is clonal in both twin1 and twin2. If a mutation is subclonal in at least one of the twins, the mutation is determined to be an EEM as it cannot be a *de novo* mutation present in the zygote. For dizygotic twins, mutations were classified as *de novo* as long as they were clonal in at least one twin.

**Mathematical framework for mutational relationships in monozygotic twins**

We developed a mathematical framework for using the estimated VAFs of monozygotic twins to infer the clonal dynamics of twin formations (**Fig. 2e**, **Supplementary Fig. S2a**). Let's assume that in the $i$ th cell division, an EEM$i$ contributes to twin1 and twin2 at $a$ and $b$ ratio, respectively. In the subsequent $i$+1 th cell division, there will be two additional daughter cells with each EEM$i$-1 and EEM$i$-2. Let's assume that EEM$i$-1 contributes to a total $c$ and $d$ fraction to Twin1 and Twin2, respectively. Since both EEM$i$-1 and EEM$i$-2 are descendants of EEM$i$, the contribution of EEM$i$-2 to each twin is determined to be $a$-$c$ and $b$-$d$, respectively, as the sum of EEM$i$-1 and EEM$i$-2's contributions in Twin1 and Twin2 should add up to $a$ and $b$. Since all $a$, $b$, $c$, and $d$ are all non-negative numbers less than 1, for a given EEM$i$'s VAF, the EEMs developed in subsequent cell divisions have to be in the left lower quadrant of EEM$i$'s VAF coordinate with EEM$i$-1's VAF coordinate as ($c$/2, $d$/2) and EEM$i$-2's coordinate as (($a$-$c$)/2, ($b$-$d$)/2) (**Supplementary Fig. S2b**). In addition, the midpoint between EEM$i$-1 and EEM$i$-2 should be ($a$/4, $b$/4) as $c$ and $d$ terms cancel out. A few example scenarios demonstrating this principle are shown in **Supplementary Fig. S2c**. For the earliest detectable cell division (i.e., $i$=1, which may be the first division from the zygote), the two mutations would be anchor mutations that would be the most helpful mutations to infer the clonal dynamics of early twinning.

**Identification of anchor mutations for clonal dynamics**

EEMs for each twin pair were used to estimate the anchor mutations (EEMs formed at the earliest detectable cell division). EEMs formed at the same cell division would be present at similar VAFs in both twins and would occur as clusters. Mutation clusters were defined as mutations where the maximum difference between mutations is less than 0.1 for both twins. For each cluster centroid coordinates were calculated. For all cluster pairs, the midpoint between each pair of centroids was calculated and the pair with closest distance to (0.25, 0.25) was chosen as the anchor mutation clusters. Anchor mutation coordinate $(a, b)$ was chosen so that $a$ and $b$ are placed with the following constraints. $0.25 \leq a \leq 0.5$, $0 \leq b \leq 0.5$, $a \geq b$, $a + b \geq 0.5$. If the best midpoint of the centroid pairs is > 0.1 away from (0.25, 0.25) or no cluster pairs are identified, only a single mutation cluster closest to (0.25, 0.25) is identified. The unidentified cluster is estimated to be present with (0.25, 0.25) as the midpoint. For each mutation cluster, if the variant had phasing information to support clonality (i.e., only two haplotypes exist), then $a$ is adjusted to 0.5 and $b$ was estimated using the same rule as above.

For monozygotic triplets, anchor mutation clusters were identified by comparing two triplet individuals at a time. EEMs that were clonal in both twins were excluded from anchor mutation calculation as these appear as *de novo* mutations due to shared common ancestor that is not the zygote.

**Calculation of genetic distances between monozygotic twins**

All EEMs identified for each twin pair were used for the calculation of the genetic distance between the two twins. For each EEM *e*, the absolute difference between the VAF in twin1 ($VAF_{1,e}$ ) and the VAF in twin2 ($VAF_{2,e}$ ) was calculated. The VAF differences between the two twins were summed and multiplied by two to get the total genetic loci differences measured in Manhattan distance.

$$Genetic\ Distance\ = 2 \sum_{e \in EEM} \left| VAF_{1,e}\ - VAF_{2,e} \right|$$

**Detection of hematopoietic mixing in the monochorionic monozygotic twins**

EEMs exclusively detected in only one of the two twins' buccal DNA sequences were used to estimate the degree of hematopoietic mixing between the twins. Reference allele and alternate allele counts in the blood sequences were used to estimate the degree of hematopoietic mixing between the two monozygotic twins using maximum likelihood estimation as described in our previous work (Yoon et al. 2022). For each EEM *e* exclusively found in only one of the twins, the expected alternative allele counts in blood with $x$ : *1-*$x$ ratio of mixing from Twin1 and Twin2 respectively can be described as

$$VAF_e^{blood} = x\, VAF_{1,e}^{buccal}\ +\ (1-x)\, VAF_{2,e}^{buccal}$$

Then, for each blood sample (Twin1 and Twin2 separately), the expected alternative allele read count at EEM $e$, $k_e$ can be estimated with the following betabinomial distribution, where $\rho$ is the dispersion factor, and shape parameters $\alpha$ and $\beta$ described as

$$\alpha_e\ =\ k_e\,(1-\rho)/\rho\ \ ,\, \beta_e\ =\ (1-k_e)\,(1-\rho)/\rho$$

$$k_e\ =\ BetaBinomial\ (n_e, \alpha, \beta)$$

We can estimate $\hat{x}$ using maximum likelihood estimate of beta-binomial log-likelihood across all informative EEMs (exclusively detected in only one of the twin's buccal DNA) as following equation where $k_e$ refers to the alternative allele read count and $n_e$ refers to the total read depth at the EEM e loci.

$$\hat{x} = arg \max_{x \in [0,1]} \sum_{e \in EEM, exclusive} log \ Pr(k_e \mid n_e, VAF_e^{blood}, \rho)$$

We used $\rho_{WGS}$= 0.0002568 since all blood samples were only sequenced with whole-genome sequencing.

**Single-cell sequencing of blood cells of monozygotic DCDA twins**

Mononuclear cells were isolated from the umbilical cord blood of two DCDA twins (ST314 and ST317) using Lymphoprep (STEMCELL Technologies) according to the manufacturer's instructions. Single mononuclear cells were deposited into individual wells of 96-well plates using the BD FACSAria II (BD Biosciences) and subjected to whole-genome amplification with the REPLI-g Advanced DNA Single Cell Kit (QIAGEN). Successful amplification was confirmed by DNA quantification and gel electrophoresis. Libraries were prepared from 92 and 90 MDA products from twins ST314 and ST317, respectively, using the TruSeq DNA Nano Kit (Illumina), followed by hybridization with probes targeting EEMs, as well as 40 heterozygous germline mutations, specific to each twin family. Sequencing was performed using the NovaSeq 6000 platform (Illumina) to achieve a mean depth of 660 for EEMs. Sequenced reads were aligned and processed as described above. Locus dropout was estimated as the proportion of heterozygous mutations with ≤ 1 read, and allelic dropout rates were estimated as the proportion of heterozygous mutations lacking both alleles with >1 read or with VAFs outside 0.01 to 0.99. Each EEM was considered present in a single cell if supported by more than one variant read with a VAF greater than 0.01. To account for allelic dropout during whole-genome amplification, EEMs with similar VAFs in bulk tissue sequencing were grouped, and an EEM group was considered present in a single cell if at least one member of the EEM group was detected. Single cells were then clustered based on the presence or absence of EEM groups, and those lacking all EEMs, and thus uninformative for lineage analysis, were excluded from the study (n=13 for both ST314 and ST317).

**Sequencing of the placenta and umbilical cords**

Placental tissues from four different quadrants and the umbilical cord were collected during delivery of each monozygotic twin. The tissues were stored in a PAXgene Tissue FIX Container (QIAGEN) for 48 to 72 hours to ensure complete fixation and were transferred to PAXgene Tissue STABILIZER (QIAGEN) for long-term storage. Next, the tissues were processed, embedded in paraffin, sectioned to a thickness of 20 to 30 μm, and stained with hematoxylin and eosin (Abcam). Trophoblast clusters were dissected under microscopic inspection using the

PALM MicroBeam (Zeiss), while Wharton's jelly was manually dissected under visual inspection. DNA was then extracted from trophoblast microdissections using the QIAamp DNA Micro Kit (QIAGEN) and from the umbilical cord using the DNeasy Blood & Tissue Kit (QIAGEN). Library preparation was performed using the NEBNext Ultra II FS DNA Library Prep Kit (NEB) for trophoblast cores and TruSeq DNA Nano Kit (Illumina) for umbilical cord, respectively. Whole-genome sequencing was conducted on the NovaSeq 6000 platform (Illumina), achieving a mean depth of 21.4 for trophoblast clusters and 33.6 for the umbilical cord, respectively. Variants in the placenta were considered to be present if present in > 1 read.

**Mutational signature analysis**

To demonstrate that EEMs and *de novo* mutation calls are genuine, we performed mutational signature analysis to evaluate whether mutational spectra of EEMs and *de novo* mutations recapitulate the mutational signature previously reported for EEMs and *de novo* mutations (Park et al. 2021; Rahbari et al. 2016). Because each sample contained only a small number of mutations per trinucleotide context, EEMs and *de novo* mutations were aggregated across samples to derive their respective mutational spectra. The mutational spectra were then decomposed by non-negative least squares method using our custom in-house script.

Each aggregated mutational spectrum was first fitted against all known signatures catalogued in COSMIC v3 (https://cancer.sanger.ac.uk/signatures/)(Alexandrov et al. 2020), which served as references for cosine similarity assessment. We next evaluated all pairwise combinations of COSMIC signatures, selected the pair with the highest cosine similarity, and then added additional signatures stepwise according to further improvements in cosine similarity. For *de novo* mutations, the fit was mostly explained by two signatures, SBS1 and SBS5, consistent with the reported signature composition of *de novo* mutations. The cosine similarity between the all-signature fit and the SBS1+SBS5 fit differed by only 0.005, with the final cosine similarity being 0.983. For EEMs, the final fit included SBS1, SBS5, SBS10b, and SBS40, yielding a cosine similarity of 0.935, only 0.025 lower than the fit with all signatures.

**Statistical analysis**

Statistical analysis was performed with R (v4.4.1). Genetic differences between twins with fetal membrane subgroups were compared using the Kruskal-Wallis test, followed by post hoc analysis with Welch's t-test to identify the statistical differences between pair-wise comparisons. The association between fetal membranes and clonal subtypes was assessed with Fisher's test.

Fisher Monte–Carlo simulation was used for the 3x3 comparison between twin fetal membranes and clonality, and Fisher's exact test was used for 2x2 comparisons. Pairwise comparisons were adjusted using Bonferroni method to control the family-wise error rate.

**Data and script availabilities**

All whole-genome, targeted sequence, and single-cell MDA amplified PBMC BAM files have been uploaded to the European Genome-Phenome Archive with accession EGAS00001005998. Custom scripts used for this project have been uploaded to https://github.com/cjyoon/mztwin

### Supplementary Figures

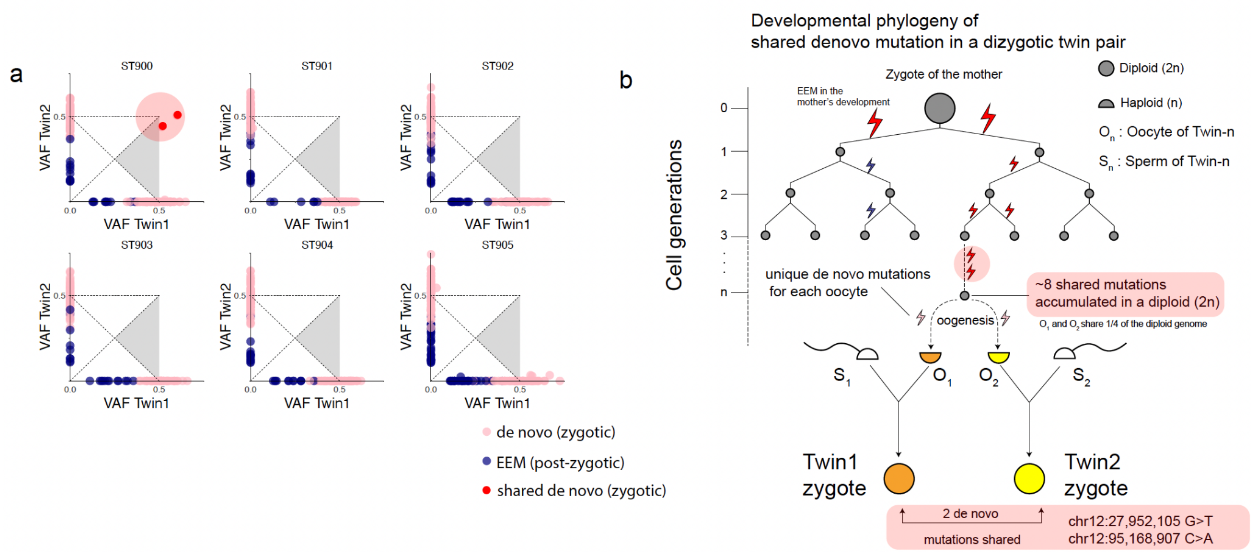


**Supplementary Figure S1. Embryonic mutation history of dizygotic twins** (a) *De novo* mutations and EEMs detected in dizygotic twin pairs. Clonal mutations are shown in pink circles. Subclonal mutations are shown in blue circles. Two shared *de novo* mutations between the dizygotic twins of ST900 are shown in red circles. (b) Embryonic history of dizygotic twin ST900 with two shared *de novo* mutations. EEMs are shown in red lightning. Haploid cells (oocytes and sperms) are shown as half circles. Diploid cells (fertilized zygotes) are shown in full circles. A large full circle at the top is the mother at the zygote stage.

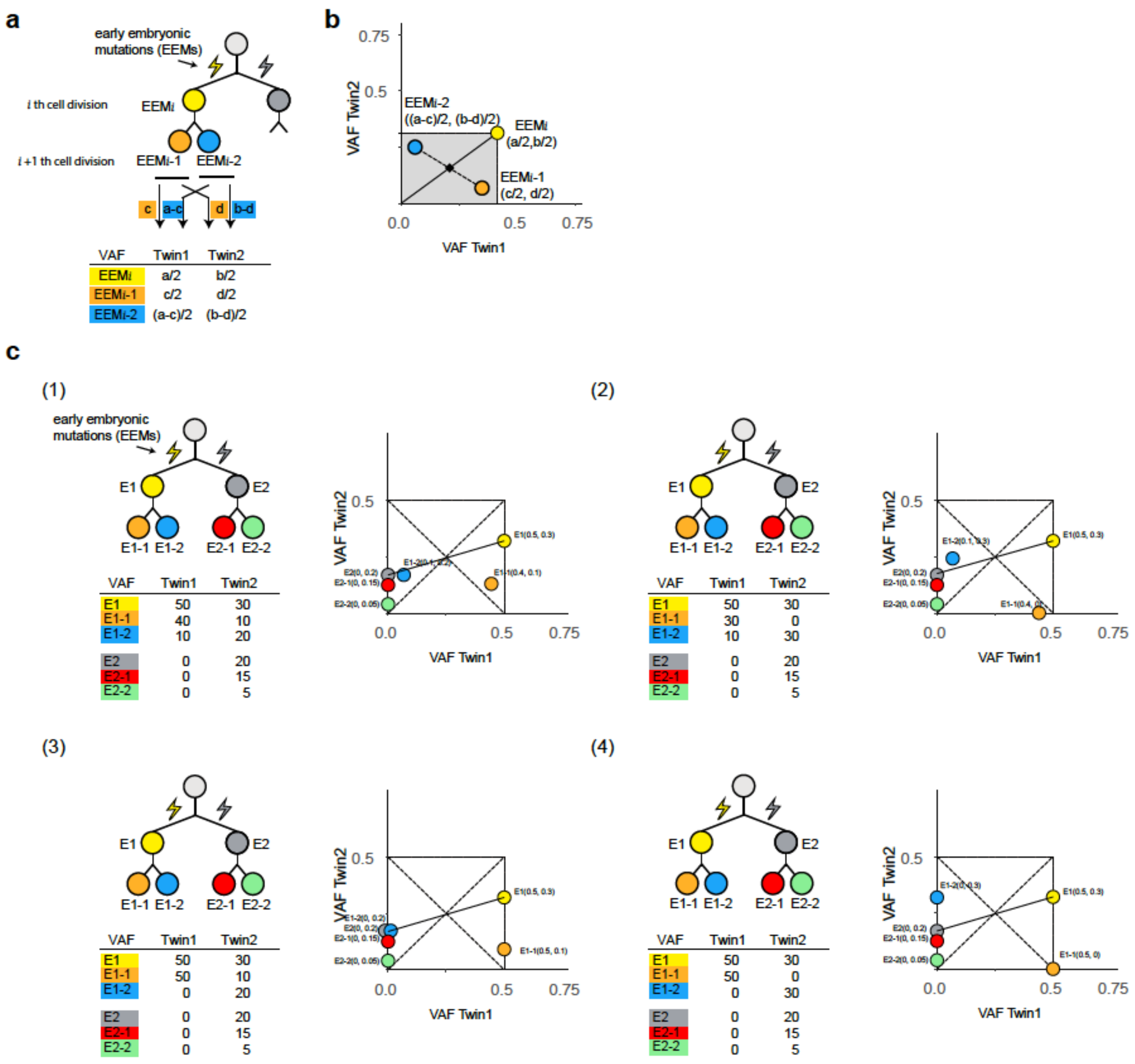


**Supplementary Figure S2. Generalized mathematical principle between EEMs in monozygotic twins.** (a) Relationships between the VAFs from EEMs are shown on a two dimensional VAF plot. Contributions of EEM*i* to twin1 and twin2 are *a* and *b*, respectively. Contributions of EEMi-1 to twin1 and twin2 are *c* and *d*, respectively. VAFs of each mutation in each twin are shown in the table. (b) VAFs of EEM*i*, EEM*i*-1, and EEMi-2 are shown on a two dimensional plot. Black dot indicates the midpoint between the origin and EEM*i* as well as the midpoint between EEM*i*-1 and EEM*i*-2. Shaded area represents the possible combination of VAFs for EEM*i*-1 and EEM*i*-2 (c) Four scenarios of different contributions of EEMs with their respective two dimensional VAF plot.

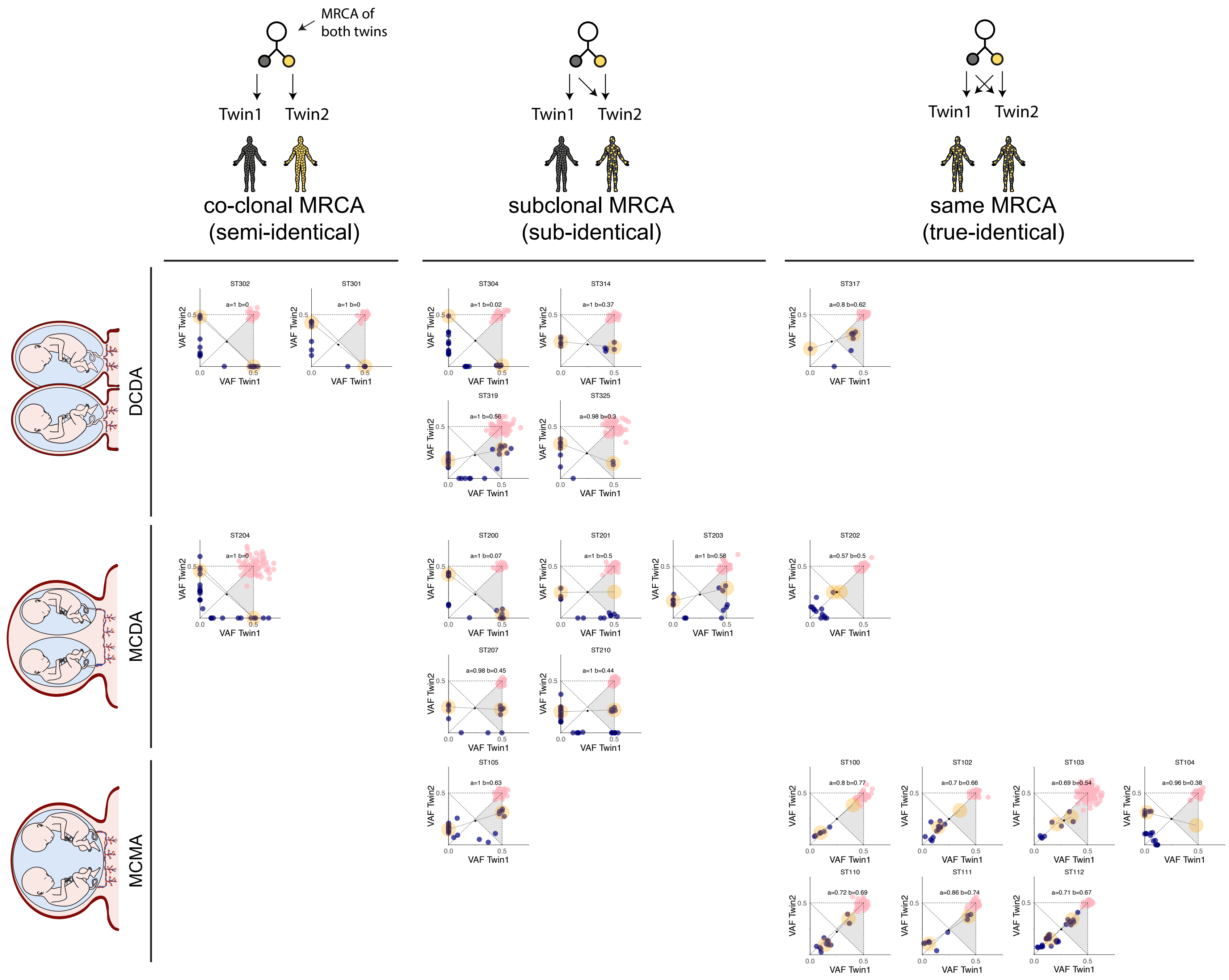

MRCA of both twins
Twin1 Twin2
co-clonal MRCA
(semi-identical)
Twin1 Twin2
subclonal MRCA
(sub-identical)
Twin1 Twin2
same MRCA
(true-identical)
DCDA
MCDA
MCMA
ST302 a=1 b=0
ST301 a=1 b=0
ST304 a=1 b=0.02
ST314 a=1 b=0.37
ST319 a=1 b=0.56
ST325 a=0.98 b=0.3
ST317 a=0.8 b=0.62
ST204 a=1 b=0
ST200 a=1 b=0.07
ST201 a=1 b=0.5
ST203 a=1 b=0.58
ST207 a=0.98 b=0.45
ST210 a=1 b=0.44
ST202 a=0.57 b=0.5
ST105 a=1 b=0.63
ST100 a=0.8 b=0.77
ST102 a=0.7 b=0.66
ST103 a=0.69 b=0.54
ST104 a=0.96 b=0.38
ST110 a=0.72 b=0.69
ST111 a=0.86 b=0.74
ST112 a=0.71 b=0.67
VAF Twin1
VAF Twin2

**Supplementary Figure S3. EEM and *de novo* mutations detected in all monozygotic twin pairs.** *De novo* mutations and EEMs are represented as pink circles and blue circles, respectively. Twins are grouped based on their fetal membranes (MCMA, MCDA, and DCDA) and their twin types, categorized by anchor mutations (full-identical, sub-identical, and para-identical). For each twin pair, the estimated *a* and *b* values are written and also marked on the two-dimensional VAF plots. Locations of the anchor cluster and counter-anchor cluster are highlighted with yellow backgrounds

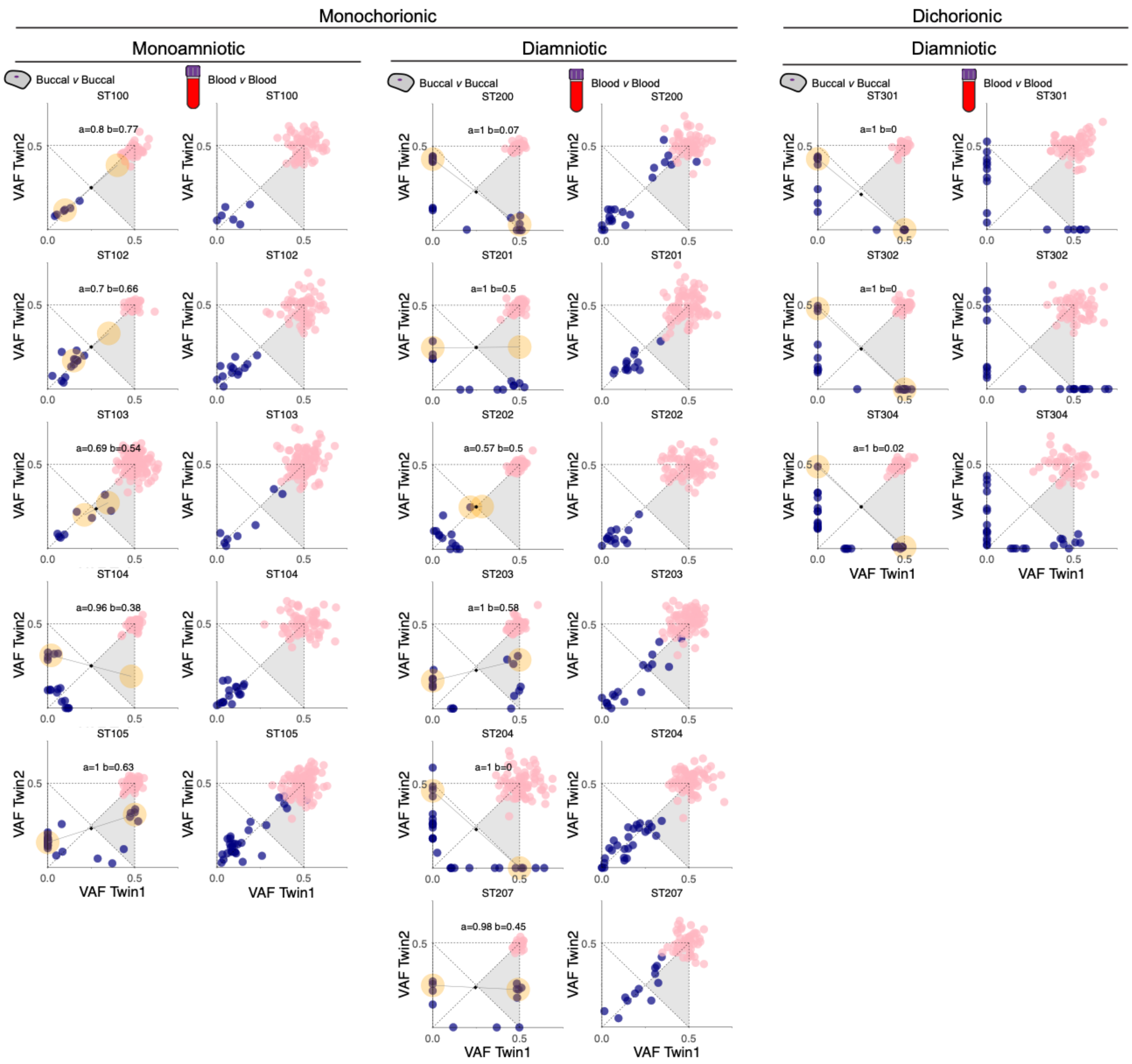


**Supplementary Fig S4. Monochorionic (MCMA and MCDA) twins share a common circulation system, and dichorionic (DCDA) twins have a separate circulation system.** Comparison of EEM and *de novo* mutations identified in 14 monozygotic twins with blood and buccal DNA. Blue circles represent EEMs and pink circles represent *de novo* mutations identified.
